# Platelet phosphatidylserine is the critical mediator of thrombosis in heparin-induced thrombocytopenia

**DOI:** 10.1101/2022.09.13.507721

**Authors:** Jan Zlamal, Anurag Singh, Karoline Weich, Hisham Jaffal, Günalp Uzun, Karina Althaus, Tamam Bakchoul

## Abstract

Heparin-induced thrombocytopenia (HIT) is a severe immune-mediated prothrombotic disorder caused by antibodies reactive to complexes of platelet factor 4 and heparin. Platelets (PLTs) and their interaction with different immune cells contribute to prothrombotic conditions in HIT. However, the exact mechanisms and the role of different PLT subpopulations to this prothrombotic enviroment remain poorly understood. In this study, we observed that HIT patient antibodies (Abs) induce relevant changes in PLT phenotype, with the key features being increased P-Selectin expression and procoagulant phosphatidylserine (PS) externalization. Formation of procoagulant PLTs was dependent on engagement of PLT Fc-gamma-RIIA by HIT Abs and resulted in significant increase of thrombin generation on the PLT surface. Using an ex vivo thrombosis model and multi-parameter assessment of thrombus formation, we observed that HIT Ab-induced procoagulant PLTs propagated formation of large PLT aggregates, leukocyte recruitment and most importantly, fibrin network generation. These prothrombotic conditions were prevented via the upregulation of PLTs intracellular cAMP with Iloprost, a clinically approved prostacyclin analogue. Additionally, the functional relevance of high P-Selectin and PS levels on procoagulant PLTs was dissected. While inhibition of P-Selectin did not affect thrombus formation, the specific blockade of PS with Lactadherin prevented HIT Ab-mediated thrombin generation and most importantly procoagulant PLT-mediated thrombus formation ex vivo. Taken together, our findings indicate that procoagulant PLTs are critical mediators of prothrombotic conditions in HIT. Upregulation of cAMP with Iloprost or PS targeting specifc therapeutics could be a promising approach to prevent thromboembolic events in HIT patients.

**Key points:**

- HIT immune complexes drive procoagulant platelet formation
- Phosphatidylserine blockade prevents HIT antibody-induced thrombus formation

## Introduction

Heparin-induced thrombocytopenia (HIT) is an immune-mediated prothrombotic disorder that is characterized by low platelet (PLT) counts and high risk of thromboembolic events.^1–3^ HIT is caused by antibodies (Abs) of the immunoglobulin G (IgG) subclass that recognize complexes of the endogenous protein platelet factor 4 (PF4) and heparin.^4,5^ These immune complexes evolve 4-14 days after heparin exposure and harbour the potential to elicit prothrombotic conditions due to their interaction with PLTs, neutrophils and monocytes via low-affinity immune receptor Fc-gamma-RIIA.^6–8^

Despite low PLT counts, HIT is associated with an increased risk of thrombosis.^9,10^ In fact, Fc-gamma-RIIA-mediated PLT activation by HIT immune complexes is associated with intense thrombin generation and prothrombotic potential.^7,11–13^ Additionally, binding of HIT Abs to PF4 and glycosaminoglycans (GAGs) on the surface of monocytes has been reported to result in increased tissue factor and microparticle release which promotes thrombin generation and PLT activation.^6,14,15^ Furthermore, HIT immune complexes can activate neutrophils via Fc-gamma-RIIA and lead to the formation of PLT-neutrophil aggregates as well as release of prothrombotic neutrophil extracellular traps (NETs).^16^ Although this evidence directs towards a prominent role of PLTs in the pathogenesis of HIT, a better understanding of HIT Ab-mediated PLT changes and the potential relevance of different PLT subpopulations in prothrombotic conditions in HIT patients is needed.

Procoagulant PLTs, a distinct subpopulation of activated PLTs, are characterized by an increased externalization of the negatively charged phospholipid phosphatidylserine (PS).^17,18^ High levels of PLT PS are efficient to promote assembly of coagulation complexes and subsequent thrombin burst.^19,20^ Recently, we showed that procoagulant PLTs might contribute to the prothrombotic state in patients with COVID-19 and vaccine induced immune thrombotic thrombocytopenia (VITT).^21,22^

In the current study, we hypothesized that anti-PF4/heparin IgG Abs from HIT patients (HIT Abs) induce a procoagulant PLT response that contributes to an increased hypercoagulable state. Our analyses revealed that the presence of heparin resulted in the formation of a HIT Ab-induced procoagulant PLT phenotype that harbours dramatic prothrombotic potential. HIT Ab-induced procoagulant PLTs were found to promote increased thrombin generation and thrombus formation in a Fc-gamma-RIIA-dependent manner that could be prevented by upregulation of PLTs’ intracellular cyclic-adenosine monophosphate (cAMP). Most importantly, HIT Ab-induced procoagulant PLT-mediated thrombin and thrombus formation were abrogated by specific inhibition of PLT PS with Lactadherin. Based on our mechanistic studies we conclude that HIT Ab-induced procoagulant PLTs might have significant relevance in prothrombotic conditions typically observed in HIT.

## Materials and Methods

### Patients and sera

Experiments were performed using serum material from HIT patients who were referred to our laboratory between March 2019 and April 2022. The diagnosis of HIT was confirmed by two experienced physicians according to current guidelines (e.g. 4Ts-score >3) as well as based on laboratory findings in enzyme-immune assay (EIA) and heparin-induced platelet activation assay (HIPA).^23^ In addition, serum samples were collected from healthy blood donors (HCs) at the Blood Donation Centre Tübingen, after a written informed consent has been obtained. Serum samples were stored at −20°C and thawed prior to experiments. To exclude unspecific effects of serum components other than Abs, all sera were heat-inactivated at 56°C for 30 min, which was followed by centrifugation at 5000g for 5 min. The supernatants were used in this study.

### Testing for anti-PF4/heparin antibodies

A commercially available IgG-EIA was used in accordance to manufacturer’s instructions (Hyphen Biomed, Neuville-sur-Oise, France). The ability of sera to activate PLTs was tested using HIPA as previously described.^24^ Additional details are available in the supplemental data.

### Preparation of washed platelets

Washed PLTs (wPLTs) were prepared from venous blood samples as previously described.^24,25^ Additional details are available in the supplemental data.

### Immunoglobulin G preparation

IgG fractions were isolated from HIT as well as from control sera using a commercially available IgG-purification-kit. Additional details are available in the supplemental data.

### Treatment of PLTs with sera/IgGs

37.5 μL of wPLTs or PRP were supplemented with 5 μL serum/IgG from HIT patients or controls and incubated for 1 hour under rotating conditions at RT. Changes in the expression of CD62p and PS were assessed via flow-cytometry (FC). Additional details are available in the supplemental data.

### Analysis of HIT antibody-induced mechanomolecular signaling mechanisms

To investigate the underlying mechanisms that result in HIT Ab-induced procoagulant PLT effects, 75 μL of wPLTs were pretreated with the Fc-gamma-RIIA blocking monoclonal antibody (moAb) anti-CD32 (moAb IV.3; Stemcell™ technologies, Vancouver, Canada) or a monoclonal isotype control (clone SC-2025; Santa Cruz Biotechnology, Dallas, USA) for 30 min at RT prior to HIT IgG treatment. To investigate the role of upregulated intracellular levels of cAMP, PLTs were pretreated with the prostacyclin analogue Iloprost (20 nM; Sigma-Aldrich, St. Louis, USA) or vehicle for 5 min at RT prior to incubation with IgG from HIT patients or HCs.

### Specific blockade of platelet surface P-Selectin and phosphatidylserine

P-selectin (CD62p) was blocked via the pretreatment of PLTs with 5 μg/mL anti-SELP humanized Ab (anti-CD62p, 15 min, RT; ProteoGenix, Schiltigheim, France). Using a modified FC approach that detects PLT-leukocyte aggregates, 5 μg/mL of anti-CD62p were found to efficiently block the interaction of TRAP-6 (10μM, 30 min, RT) activated wPLTs with neutrophil granulocytes that were purified via density gradient centrifugation (data not shown).^26–28^

Specific inhibition of externalized PS on the surface of procoagulant PLTs was performed via the incubation of PLTs with Lactadherin, a calcium independent ligand of PS (200nM, 15 min, RT; Haematologic Technologies, Essex Junction, USA).^29,30^ Sufficient blocking concentrations of Lactadherin were assessed via Calibrated Automated Thrombogram (CAT; data not shown).

### Thrombin generation assay

HIT Ab-induced thrombin generation on PRP was detected using CAT (Stago, Maastricht, Netherlands) according to the manufacturer’s instructions. Additional details are available in the supplemental data.

### Investigation of antibody-induced thrombus formation

To investigate the effect of Ab-induced PLT alterations on the coagulation cascade and subsequent thrombus formation, an ex vivo model for thrombus formation (BioFlux 200, Fluxion Biosciences, Alameda, USA) was used according to the recommendations of the ISTH Standardization Committee for Biorheology.^31^ Briefly, microfluidic channels were coated with collagen (100 μg/mL; Collagen Horm, Takeda, Linz, Austria) overnight at 4°C and blocked with 1% of human serum albumin (Kedrion, Barga, Italy) 1 h at RT before perfusion. WB of healthy individuals with blood group O was collected into 0.105 M sodium citrate containing monovettes (Sarstedt, Nuembrecht, Germany) that were supplemented with Corn trypsin inhibitor (5 μg/mL; Santa Cruz Biotechnology, Dallas, USA) and allowed to rest for 30 min at RT. After splitting the WB into aliquots of 200 μL, PRP was isolated via centrifugation (15 min, 120g, RT, no break). Afterwards, 37.5 μL of the supernatant PRP were gently separated and incubated with 5 μL control or HIT IgG under rotating conditions (60 min, RT). After the incubation period, samples were labelled with 3,3’-Dihexyl-oxacarbocyaniniodid (DiOC_6_, 2.5 μM; Sigma Aldrich, Saint Louis, USA), Alexa Fluor (AF) 647-Annexin A5 (1:200), AF 546 human fibrinogen (8.5 μg/mL) and Hoechst 33342 (3 μg/mL; Thermo Scientifc, Carlsbad, USA) during reconstitution into WB. When indicated, the separated PRP was pretreated with moAb IV.3 or isotype control at a concentration of 20 μg/mL for 30 min at RT. For cAMP elevation in PLTs, PRP was pretreated with Iloprost (20 nM) or vehicle control (5 min, RT) prior to IgG incubation. For specific inhibition of CD62p and PS, PLTs were incubated with anti-CD62p Ab (5 μg/mL) and Lactadherin (200nM) for 15 min at RT. To avoid excess amounts of Lactadherin that could interfere with different blood coagulation factors, PLTs were washed (700g, RT, no brake) and resuspended with fresh PLT poor plasma after HIT IgG incubation was performed. Finally, reconstituted WB samples were recalcified and run at a shear rate of 250s^-1^ (10 dyne) for 10 min as previously described.^25,32^

### Image acquisition and analysis

Immunofluorescence images were aquired from 3-5 randomly chosen microscopic fields in different fluorescence channels (x40 magnification) using a Zeiss Axio Observer 7 (Carl Zeiss, Oberkochen, Germany). Images were processed identically using adjusted threshold settings and exclusion of image artefacts by using Fiji image processing software.^33^ Thrombus formation was determined by measuring the surface area coverage (SAC) by AF647 Annexin-V, DiOC_6_ and AF546 human-Fibrin(-ogen) positive labelled structures and normalized as total (%SAC) to the whole channel area. The number of adherent leukocytes was calculated by counting the total number of Hoechst 33342 positive labelled cells of corresponding microscopic fields.

### Statistical analysis

Statistical analysis was performed using GraphPad Prism, Version 8.0 (GraphPad, La Jolla, USA). Comparison between groups was performed by Mann-Whitney *U* test for unpaired data sets and Wilcoxon matched-pairs signed-rank or paired sample t test for paired data sets. *P* values < 0.05 were considered statistically significant.

### Ethics

Studies involving human material were approved by the Ethics Committee of the Medical Faculty, Eberhard-Karls University, Tübingen, Germany, and were conducted in accordance with the Declaration of Helsinki.

## Results

### Patient characteristics

In this study, we used serum samples from 37 patients with suspected HIT. 23/37 (62%) patients were male and the mean age was 65 years (range: 37-87 years). 8/37 (22%) patients presented with low 4Ts-score (<3), 16/37 (43%) had intermediate 4Ts-score (4-5) and 13/37 (35%) show high 4Ts-score (6-8). 28/37 (76%) patients tested positive for anti-PF4/heparin IgG Abs in EIA (mean OD: 2.12, range: 0.51-3.61). The diagnosis of HIT was confirmed in 20/37 (54%) using HIPA (median time to platelet aggregation in the presence of 0.2 IU/mL heparin: 5 min, range: 5 to 25 min). In the HIT-positive cohort, thrombocytopenia (PLT count <150×10^6^/mL) was detected in 19/20 (95%) and clinical overt thrombosis was observed in 9/20 (45%) patients including 6 pulmonary embolisms and 2 occlusions in the arterial vascular bed (for more details see Table 1).

**Table 1.** Patient characteristics.

| Patient ID | gender | age | PLT-count<br>† | Mean OD<br>EIA | HIPA | Time to aggregation<br>in HIPA | Thrombosis |
| --- | --- | --- | --- | --- | --- | --- | --- |
| 1 | m | 54 | 114 | 0.109 | neg. | x | yes |
| 2 | f | 79 | 120 | 0.131 | neg. | x | no |
| 3 | f | 56 | 79 | 0.071 | neg. | x | no |
| 4 | m | 79 | 283 | 0.091 | neg. | x | no |
| 5 | m | 70 | 41 | 0.111 | neg. | x | no |
| 6 | f | 48 | 133 | 0.099 | neg. | x | no |
| 7 | m | 45 | 27 | 0.072 | neg. | x | no |
| 8 | m | 63 | 32 | 0.183 | neg. | x | no |
| 9 | f | 74 | 95 | 0.073 | neg. | x | no |
| 10 | f | 68 | 83 | 2,360 | neg. | x | yes |
| 11 | m | 67 | 424 | 1,189 | neg. | x | no |
| 12 | m | 40 | 100 | 1,776 | neg. | x | yes |
| 13 | f | 83 | 114 | 0,566 | neg. | x | no |
| 14 | m | 59 | 80 | 0,732 | neg. | x | no |
| 15 | m | 61 | 211 | 0,546 | neg. | x | yes |
| 16 | f | 67 | 103 | 0,508 | neg. | x | no |
| 17 | m | 65 | 101 | 0,728 | neg. | x | no |
| 18 | m | 76 | 142 | 2,729 | pos. | 20 | no |
| 19 | m | 71 | 59 | 2,807 | pos. | 5 | no |
| 20 | m | 77 | 184 | 2,216 | pos. | 5 | yes |
| 21 | m | 83 | 119 | 1,130 | pos. | 25 | no |
| 22 | m | 71 | 114 | 2,731 | pos. | 15 | no |
| 23 | m | 57 | 60 | 2,490 | pos. | 10 | no |
| 24 | f | 79 | 98 | 1,740 | pos. | 10 | no |
| 25 | f | 74 | 35 | 1,967 | pos. | 10 | no |
| 26 | f | 41 | 135 | 2,205 | pos. | 10 | no |
| 27 | m | 80 | 80 | 3,190 | pos. | 5 | no |
| 28 | f | 79 | 22 | 2,871 | pos. | 5 | no |
| 29 | f | 58 | 76 | 2,755 | pos. | 15 | yes |
| 30 | m | 45 | 37 | 2,742 | pos. | 5 | yes |
| 31 | m | 77 | 63 | 2,520 | pos. | 5 | yes |
| 32 | f | 87 | 81 | 2,342 | pos. | 5 | yes |
| 33 | f | 46 | 62 | 1,864 | pos. | 5 | yes |
| 34 | m | 49 | 16 | 3,130 | pos. | 5 | no |
| 35 | m | 77 | 46 | 3,162 | pos. | 5 | yes |
| 36 | m | 37 | 20 | 2,786 | pos. | 5 | yes |
| 37 | m | 54 | 23 | 3,609 | pos. | 5 | yes |

PLT, platelet; m, male; f, female; OD, optical density; EIA, enzyme-immune assay; HIPA, heparin-induced platelet activation assay; † reference range PLT-count (150-450 x10<sup>9</sup>/L); #1-17: patients with suspected HIT; #18-37: patients with confirmed HIT.

### Platelets membrane phospholipid symmetry is rearranged by sera from HIT patients

In a first screening, wPLTs from healthy individuals were incubated with sera from 37 patients with suspected HIT in the presence of buffer, low- (0.2 IU/mL) or high- (100 IU/mL) dose heparin. As shown in Figure 1A, marked changes in CD62p expression and PS surface externalization were induced by sera from HIT patients in the presence of low-dose (0.2 IU/mL) heparin but not in the presence of high-dose (100 IU/mL) heparin (mean percent of CD62p and PS double positive PLTs [mean %]±SEM: 38.65±5.79% vs. 1.14±0.26%, p value <0.0001; Figure 1A). Contrary, no significant alterations in the expression levels of PLT CD62p and PS were observed in sera from patients with suspected but not confirmed HIT (HIPA negative) compared to sera from HCs (mean %±SEM: 0.63±0.12% vs. 1±0%, p value 0.0565; Figure 1A). To investigate whether our findings were solely due to the interaction of HIT Abs with PLTs and not induced by unspecific serum-mediated effects, we next investigated IgG fractions from corresponding HIT patients. HIT IgG isolates induced significant procoagulant PLT formation in a heparin-dependent manner compared to buffer and HC IgG (mean % ±SEM: 33.71±3.88% vs. 4.30±1.13%, p value 0.0039; and vs. 1.40±0.84%, p value 0.0091, respectively; Figure 1B).

**Figure 1.**
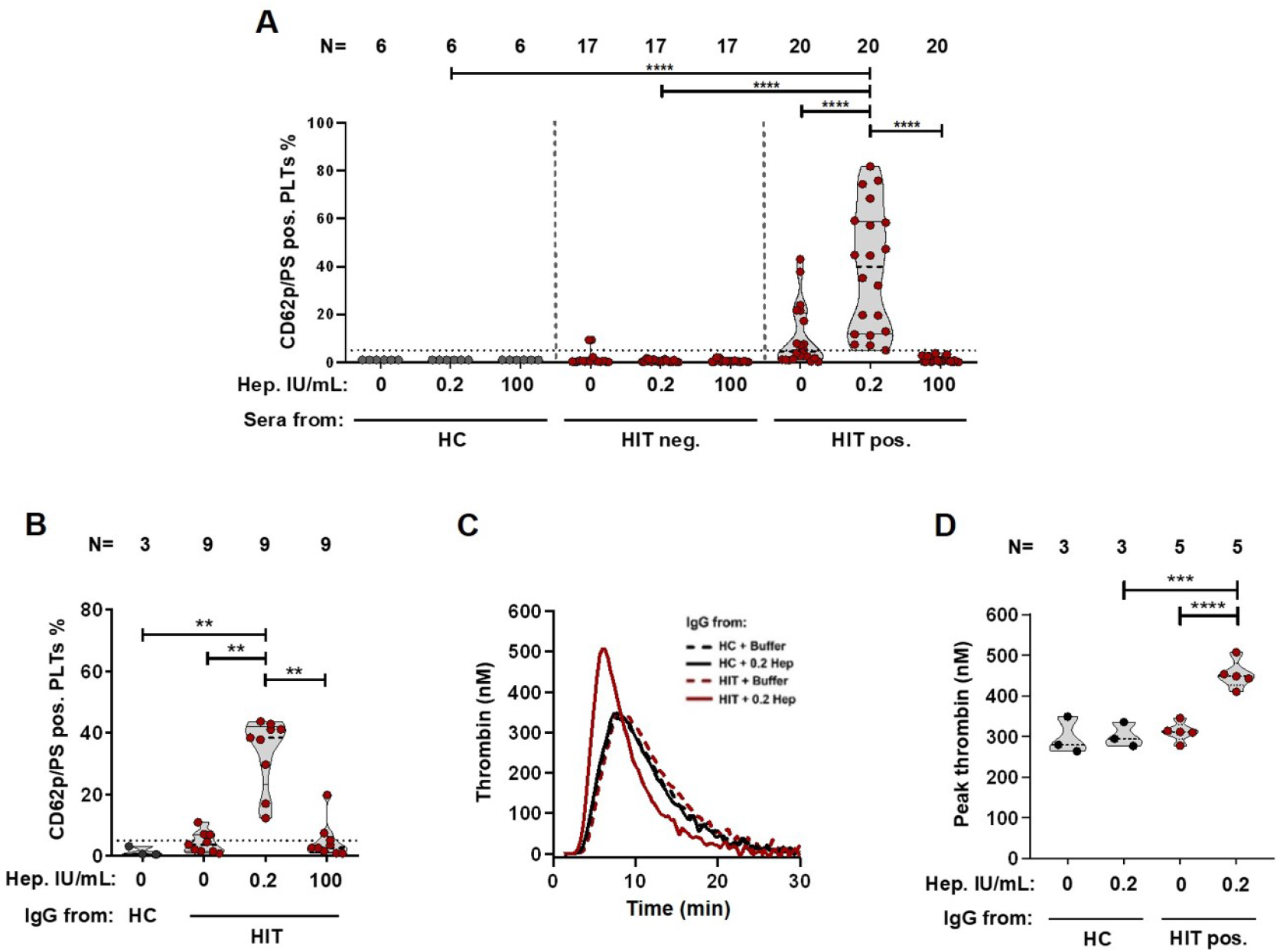
HIT antibodies induce procoagulant platelets and thrombin generation in a heparin-dependent manner. [**A**] PLTs were incubated with sera from patients with suspected HIT (n=37), control sera (HC, n=6) or [**B**] with corresponding IgG isolates and tested for changes in the expression levels of CD62p and PS via double staining in FC. Where indicated, PLTs were co-incubated with low- (0.2 IU/mL) or high- (100 IU/mL) dose heparin. [**C**] Representative thrombin generation curve induced on PLTs after incubation with IgGs from HC (black lines) or HIT patients (red lines) in the presence of buffer or heparin (0.2 IU/mL). Each curve represents the amounts of generated thrombin over time in the presence of buffer (dashed lines) or heparin (0.2 IU/mL, [solid lines]). [**D**] Data were quantified as peak thrombin generated (nM) using Thrombinoscope software and Graphpad prism. The number of patients/IgG tested is reported in each graph. Violin plots showing the distribution of the values were generated using Graphpad Prism 8. *P < 0.05, **P < 0.01, ***P < 0.001, and ****P < 0.0001. ns, non-significant; CAT, calibrated automated thrombogram; PLT, platelet; FC, flow cytometry; HC, healthy control; IgG, immunoglobulin G; CD62p, P-selectin; PS, phosphatidylserine; N, number of samples.

### HIT antibodies mediate increased thrombin generation and thrombus formation

Higher levels of thrombin were generated on PLTs upon HIT IgG incubation in the presence of low-dose (0.2 IU/mL) heparin compared to buffer and HCs (mean peak thrombin [nM]±SEM: 452.70±15.67 vs. 312.00±10.76, p value <0.0001; and vs. 297.90±26.20, p value 0.0009, respectively; Figure 1C+D).

Next, we established an ex vivo HIT-thrombosis model that utilizes tetra staining to visualize the contribution of procoagulant PLTs (Annexin-V positive, [red]), non-procoagulant PLTs (DiOC_6_ positive, [green]), leukocytes (Hoechst 33342 positive, [blue]) and fibrin (magenta) to HIT Ab-induced thrombus formation (Supplemental Figure 1). In a first proof of concept experiment, we tested whether our thrombosis model resembles the pathophysiology of HIT, namely heparin-dependent PLT activation and subsequent vessel occlusion. As shown in Figure 2, the presence of heparin (0.2 IU/mL) resulted in a significant increase of thrombus formation while such changes were not detectable under buffer conditions. Interestingly, procoagulant PLTs were observed to circulate and attach to the collagen surface of the microfluidic system from early on in our experiments (mean %SAC Annexin-V±SEM: 3.57±0.29% vs. 0.72±0.19%, p value 0.0012; Figure 2 & Supplemental Movie 1). Later, procoagulant PLTs incorporated into the growing thrombus (Supplemental Movie 2). Additionally, heparin-dependent deposition of non-procoagulant PLTs was observed in the presence of HIT Abs (mean %SAC DiOC_6_±SEM: 2.21±1.48% vs. 14.69±3.86%, p value 0.020; Figure 2 & Supplemental Movie 3). Moreover, the multicellular composition of HIT Ab-mediated thrombosis in our microfluidic system was confirmed by an increased heparin-dependent recruitment of leukocytes to the thrombi (mean no. of Hoechst positive cells±SEM: 20±13 vs. 2±0, p value 0.141; Figure 2). Finally, HIT Abs were able to activate the plasmatic coagulation system in our assays as indicated by increased fibrin network generation (mean %SAC Fibrin(-ogen)±SEM: 8.93±1.09% vs. 1.38±0.54%, p value 0.0047; Figure 2, Supplemental Movie 4). No increased thrombus formation was induced by IgGs from HCs despite the presence of low-dose (0.2 IU/mL) heparin (Supplemental Figure 2).

**Figure 2.**
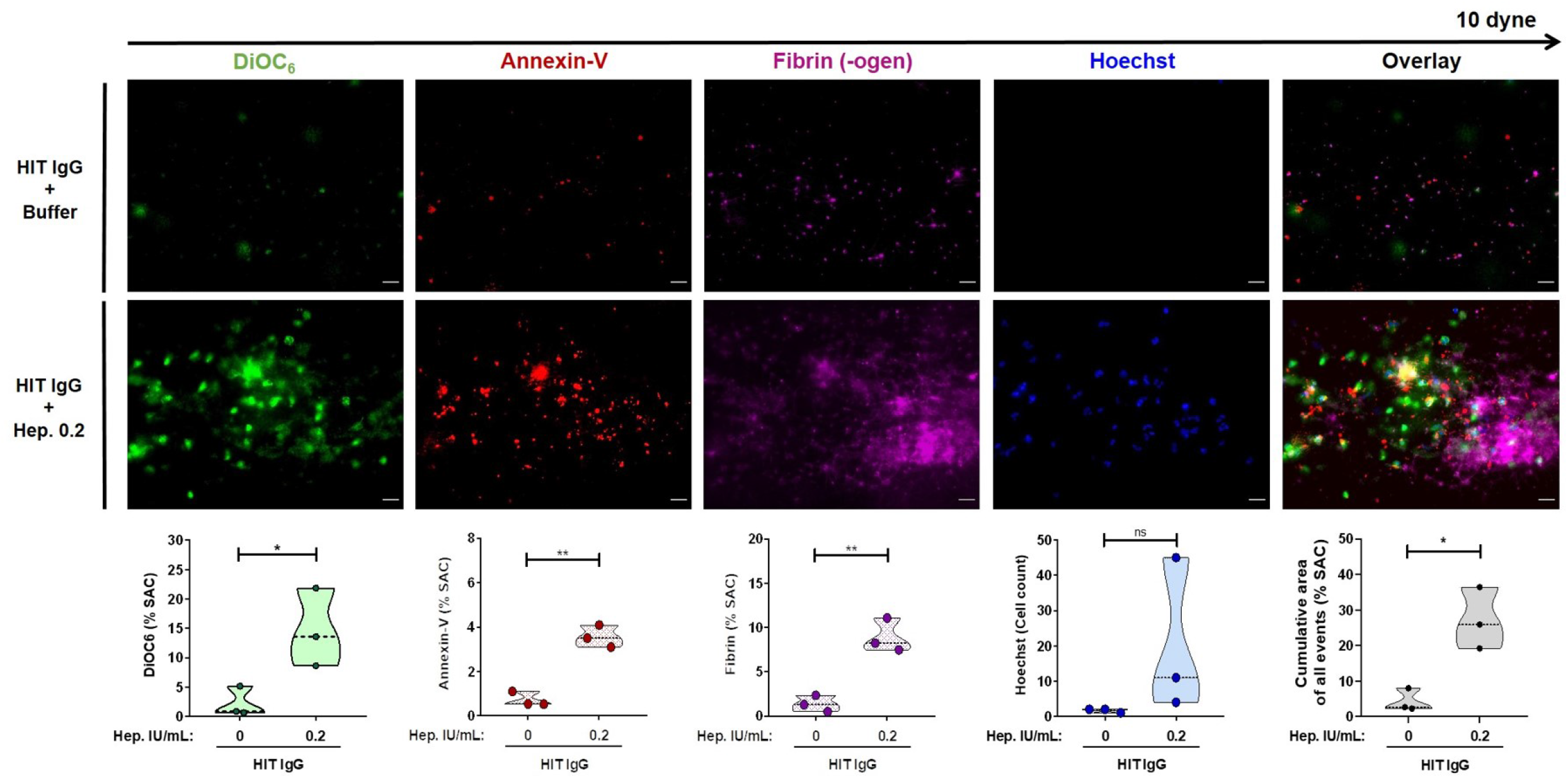
Heparin dependency of the ex vivo HIT thrombosis model. PRP from healthy individuals was incubated with IgGs from HIT patients in the presence of buffer (upper panel) or heparin (lower panel) prior to labelling of PLTs with DiOC_6_ (green), procoagulant PLTs with AF647 Annexin-V (red), AF546 Fibrinogen (magenta) and leukocytes with Hoechst 33342 (blue). After labelling, samples were reconstituted into autologous WB. Samples were then recalcified and perfused through microfluidic channels at a venous shear rate of 250s^-1^ (10 dyne) for 10 min. Images were acquired at x40 magnification in different fluorescence channels using a Zeiss Axio Observer 7 microscope. Scale bar 20μm. Images were processed identically using adjusted threshold settings and exclusion of image artefacts using Fiji image processing software. Violin plots showing the percentage of total surface area coverage (%SAC) by DiOC_6_, PS, Fibrin (-ogen), number of Hoechst positive labelled cells and cumulative area with DiOC_6_, PS and Fibrin(-ogen) labelled thrombus captured in the microfluidic channel. *p<0.05, **p<0.01 and ***p<0.001. ns, non-significant; IgG, immunoglobulin G; PLT, platelet; PRP, platelet-rich plasma, PS, phosphatidylserine.

### Fc-gamma-RIIA signaling induces procoagulant platelets in HIT

We next aimed to dissect the underlying mechano-molecular signaling pathways leading to HIT Ab-induced procoagulant PLT formation and increased prothrombotic potential. Pretreatment of PLTs with Fc-gamma-RIIA blocking moAb IV.3 significantly reduced HIT Ab-induced procoagulant PLT formation (mean %±SEM: 46.74±4.97% vs. 5.48±1.12%, p value 0.0313; Supplemental Figure 3A) as well as thrombin generation on PLTs (mean peak thrombin [nM]±SEM: 423.70±18.92 vs. 283.3±24.95, p value 0.0085; Supplemental Figure 3B+-C). Most importantly, inhibition of PLT Fc-gamma-RIIA significantly inhibited HIT Ab-induced procoagulant PLT deposition (mean %SAC Annexin-V±SEM: 4.83±0.23% vs. 1.12±0.43%, p value 0.0042; Figure 3). Similar, a significant reduction in the deposition of non-procoagulant PLTs was observed in the presence of moAb IV.3 (mean %SAC DiOC_6_±SEM: 20.99±0.97% vs. 2.69±1.15%, p value 0.0062). Additionally, PLT Fc-gamma-RIIA blockade resulted in a marked inhibition of fibrin generation and attachment of leukocytes to the thrombus surface compared to isotype control (mean %SAC Fibrin(-ogen)±SEM: 13.90±1.59% vs. 2.95±1.49%, p value 0.0317; and mean no. of Hoechst positive cells±SEM: 19±4 vs. 5±1, p value 0.0257, respectively; Figure 3).

**Figure 3.**
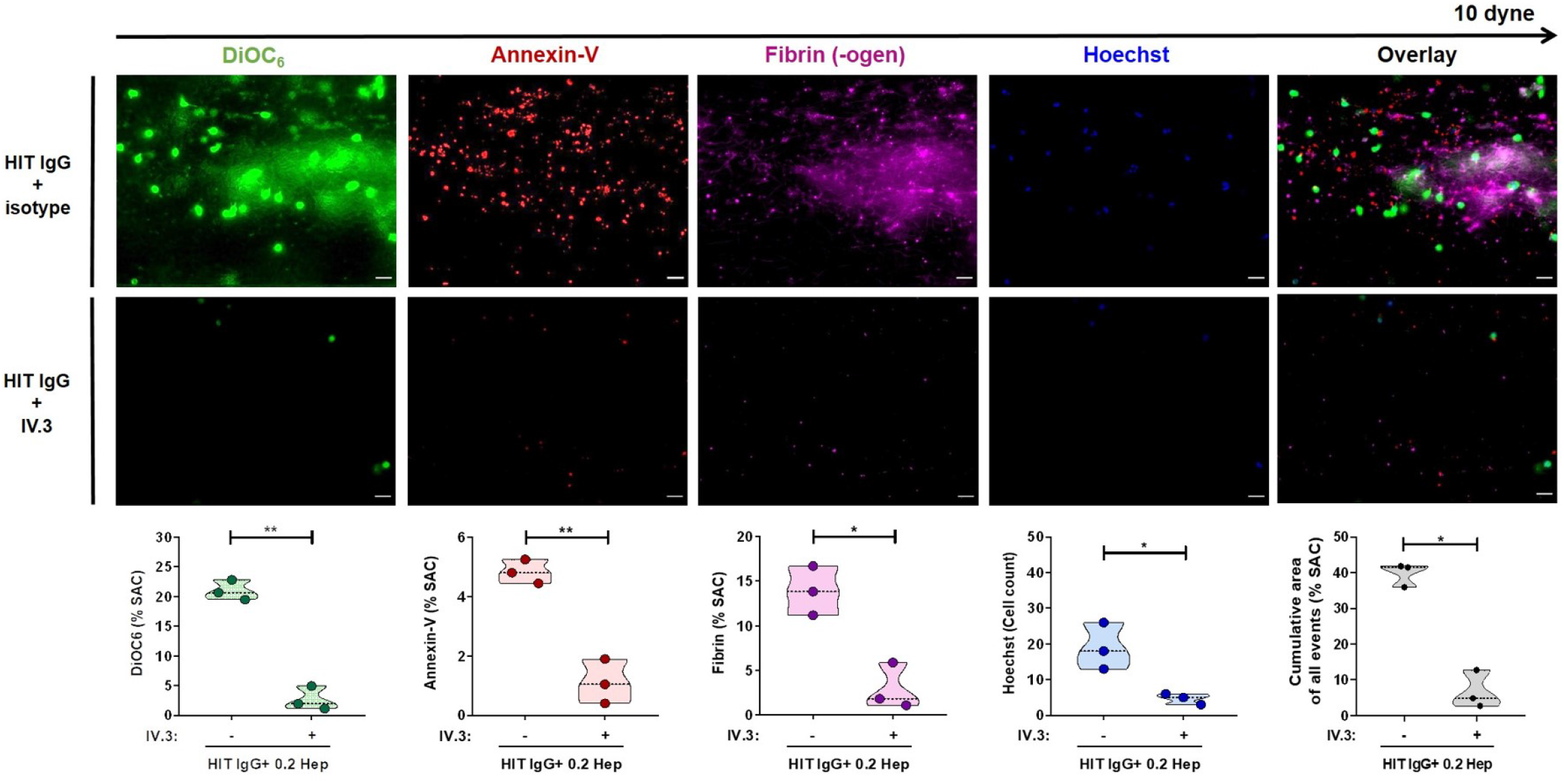
Fc-gamma-RIIA inhibition prevents thrombus formation by HIT IgG. PLTs from healthy individuals were incubated with IgG from HIT patients in the presence of heparin (0.2 IU/mL), and moAb IV.3 (lower panel) or isotype control (upper panel) before reconstitution into whole blood and perfusion through microfluidic channels at a venous rate of 250s^-1^ (10 dyne) for 10 min. After perfusion, images were acquired at x40 magnification. Scale bar 20μm. Violin plots showing the percentage of total surface area coverage (%SAC) by DiOC_6_, PS, Fibrin(-ogen), count of Hoechst positive labelled cells and cumulative %SAC with DiOC_6_, PS and Fibrin(-ogen) labelled thrombus captured in the microfluidic channel. *p<0.05, **p<0.01 and ***p<0.001. PLT, platelet; IgG, immunoglobulin G; moAb, monoclonal antibody; PS, phosphatidylserine.

### Antibody-induced thrombosis in HIT is mediated by calcium dependent signaling pathways

Sustained high levels of intracellular calcium are a typical feature of procoagulant PLTs.^34^ Prostacyclins are well characterized to inhibit intracellular calcium release via binding to PLTs prostacyclin receptor (IP-R), activation of membrane bound adenylyl cyclase and subsequent elevation of intracellular cAMP.^35^ The pretreatment of PLTs with the prostacyclin analogue Iloprost clearly showed the potential to inhibit HIT Ab-induced procoagulant PLT formation (mean %±SEM: 35.99±5.98% vs. 4.20±1.08%, p value 0.0313; Supplemental Figure 4A). This inhibition was not only observed phenotypically as Iloprost pretreatment also prevented from procoagulant PLT-induced increased thrombin generation (mean peak thrombin [nM]±SEM: 295.80±17.70 vs. 125.80±7.08, p value 0.0026; Supplemental Figure 4B+C). Most importantly, no elevated deposition of procoagulant PLTs was observed when PLTs were pretreated with Iloprost prior to HIT IgG incubation (mean %SAC Annexin-V±SEM: 4.09±0.24% vs. 1.42±0.19%, p value 0.0003; Figure 4). Similarly, the attachment of non-procoagulant PLTs was almost completely inhibited with Iloprost compared to the vehicle control (mean %SAC DiOC_6_±SEM: 17.97±2.25% vs. 2.19±0.76%, p value 0.0054). Additionally, Iloprost pretreatment reduced fibrin deposition and prevented the recruitment of leukocytes to thrombi (mean %SAC Fibrin(-ogen)±SEM: 10.40±1.41% vs. 3.49±0.90%, p value 0.0037; and mean no. of Hoechst positive cells±SEM: 14±5 vs. 2±0, p value 0.0257, respectively; Figure 4).

**Figure 4.**
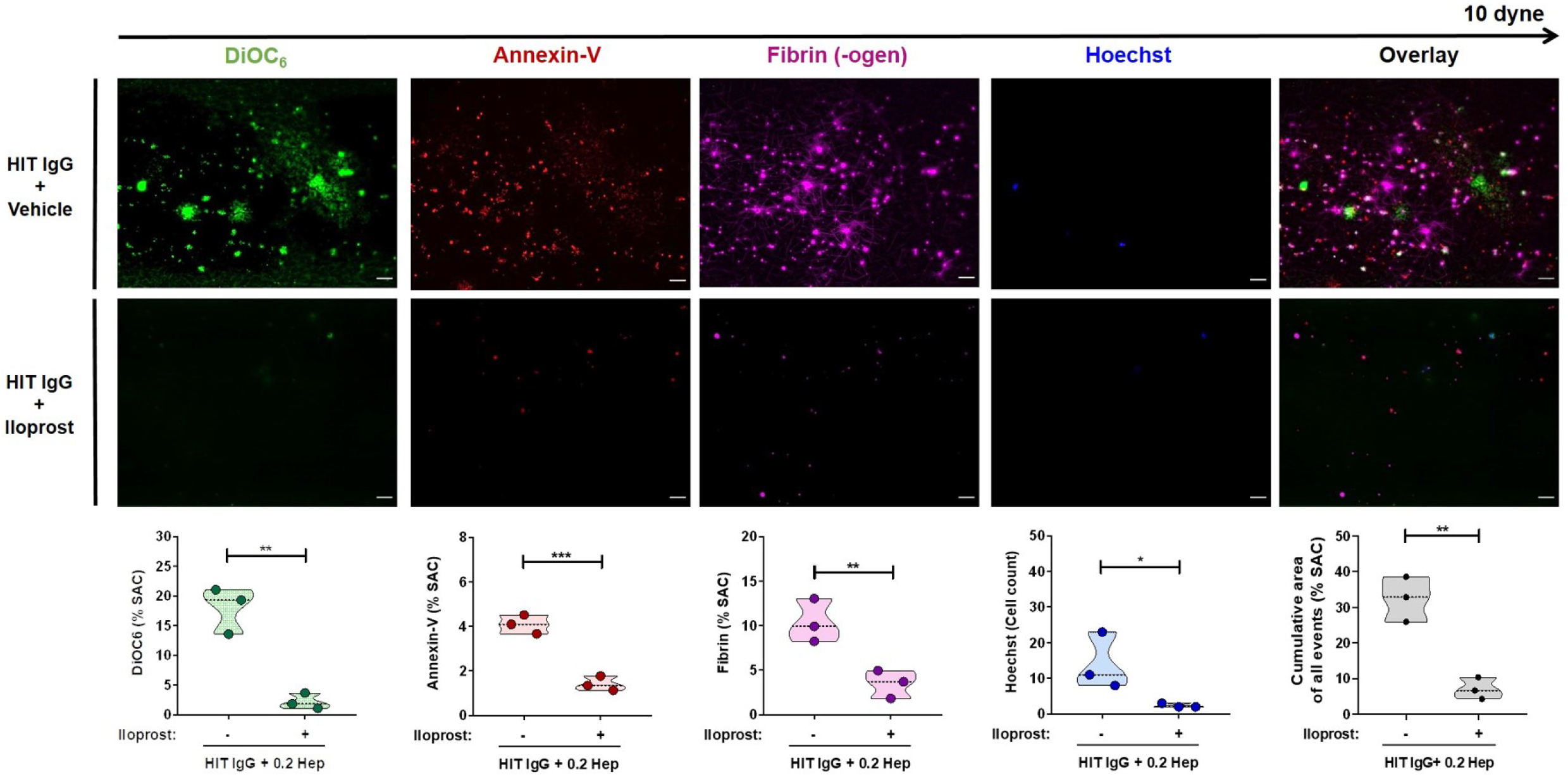
Upregulation of platelet cAMP protects from HIT Ab-induced thrombus formation. PLTs from healthy individuals were incubated with IgG from HIT patients in the presence of vehicle (upper panel) or Iloprost (20 nM, [lower panel]) and heparin (0.2 IU/mL). After reconstitution into autologous whole blood and recalcification, samples were perfused through microfluidic channels at a venous shear rate of 250s^-1^ (10 dyne) for 10 min. After perfusion, images were acquired at x40 magnification. Scale bar 20μm. Violin plots showing the percentage of total surface area coverage (%SAC) by DiOC_6_, PS, Fibrin (-ogen), count of Hoechst positive labelled cells and cumulative %SAC with DiOC_6_, PS and Fibrin (-ogen) labelled thrombus captured in the microfluidic channel. *p<0.05, **p<0.01 and ***p<0.001. PLT, platelet; IgG, immunoglobulin G; PS, phosphatidylserine.

### Phosphatidylserine but not P-Selectin is essential to propagate HIT antibody-mediated thrombosis

HIT Ab-induced procoagulant PLT formation and subsequent prothrombotic changes were significantly inhibited via PLT Fc-gamma-RIIA blockade as well as intracellular cAMP elevation. However, these data do not allow a conclusion, whether Ab-induced procoagulant PLTs are the causing factor for an increased prothrombotic potential.

We observed a slight inhibition of thrombin generation with anti-CD62p Ab, both in HC and HIT IgG incubated PLTs. However, HIT IgGs were able to induce significant increase in thrombin generation on the procoagulant PLT surface compared to HC (Figure 5A+B). In our ex vivo thrombosis model, anti-CD62p Ab was able to reduce leukocyte deposition on the thrombus surface (mean no. of Hoechst positive cells±SEM: 16±3 vs. 4±1, p value 0.0434; Figure 6). However, this reduction of leukocyte recruitment did not seem to affect thrombus formation as no significant alterations in the deposition of procoagulant and non-procoagulant PLTs was observed (mean %SAC Annexin-V±SEM: 4.80±0.67% vs. 5.27±0.21%, p value 0.262; and mean %SAC DiOC_6_±SEM: 17.85±2.91% vs. 19.78±1.78%, p value 0.124, respectively). Moreover, no changes in plasmatic coagulation, namely fibrin generation, were observed in the presence of anti-CD62p (mean %SAC Fibrin(-ogen)±SEM: 9.71±1.42% vs. 11.68±2.31%, respectively, p value 0.256; Figure 6).

**Figure 5.**
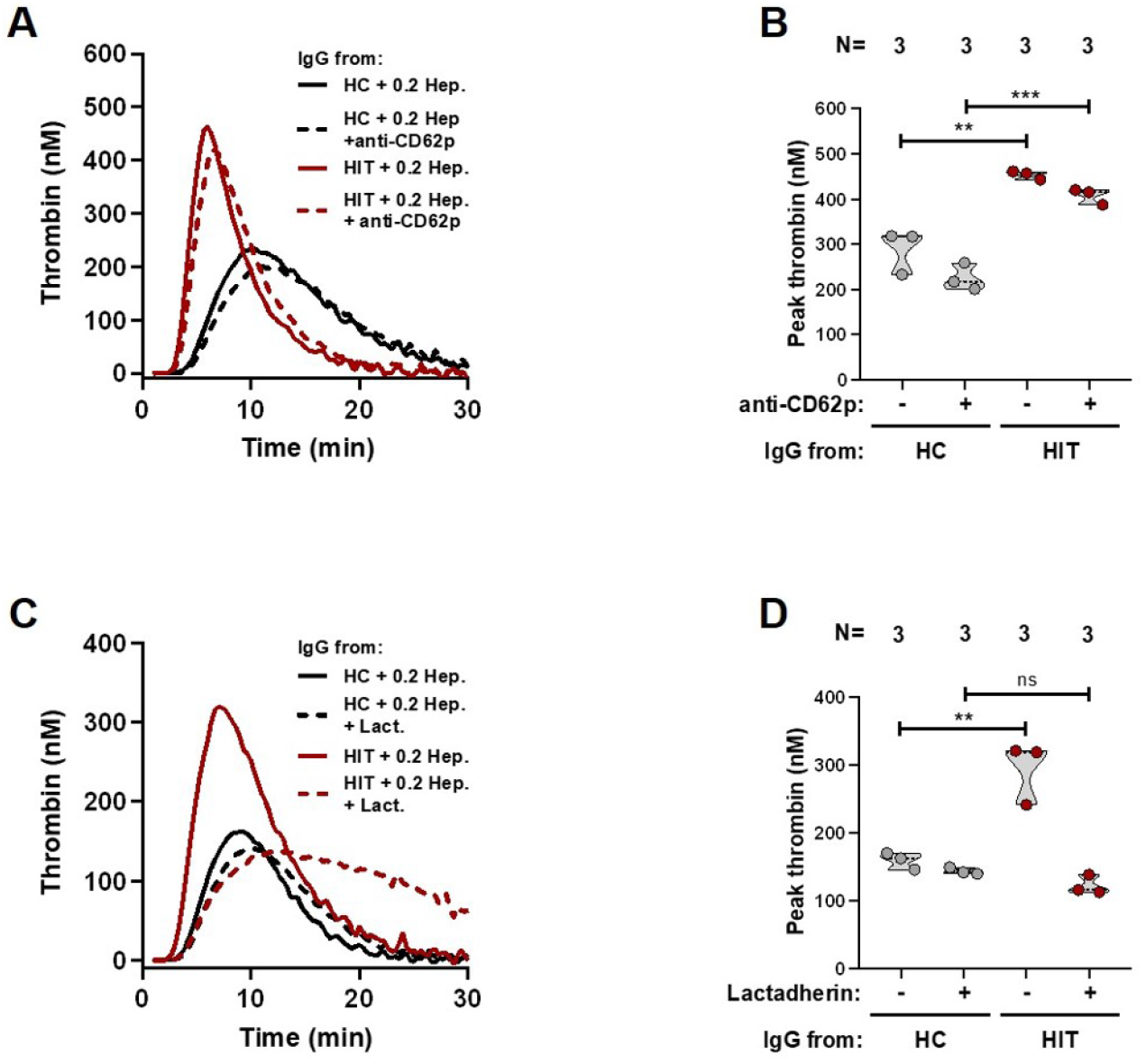
Increased platelet phosphatidylserine causes higher thrombin generation in HIT. Panel [**A**] shows thrombin generation potential on PLTs after incubation with IgGs from HIT patients (red line) or HC (black line) in the presence of vehicle or anti-CD62p blocking Ab. Panel [**C**], thrombin generation on PLTs that were incubated with different HIT patient IgGs (red line) or HC IgG (black line) and treated with Lactadherin or vehicle, before CAT analysis was performed. Each curve represents the amounts of generated thrombin over time induced by HIT IgG in the presence of vehicle (solid lines) or anti-CD62p blocking Ab or Lactadherin (dashed lines), respectively. [**B+D**] Data were quantified as peak thrombin generated (nM) using Thrombinoscope software and Graphpad prism. *p<0.05, **p<0.01 and ***p<0.001. ns, non-significant; CAT, calibrated automated thrombogram; CD62p, P-Selectin; PLT, platelet; HC, healthy control; Ab, antibody; IgG, immunoglobulin G; N, number of samples.

**Figure 6.**
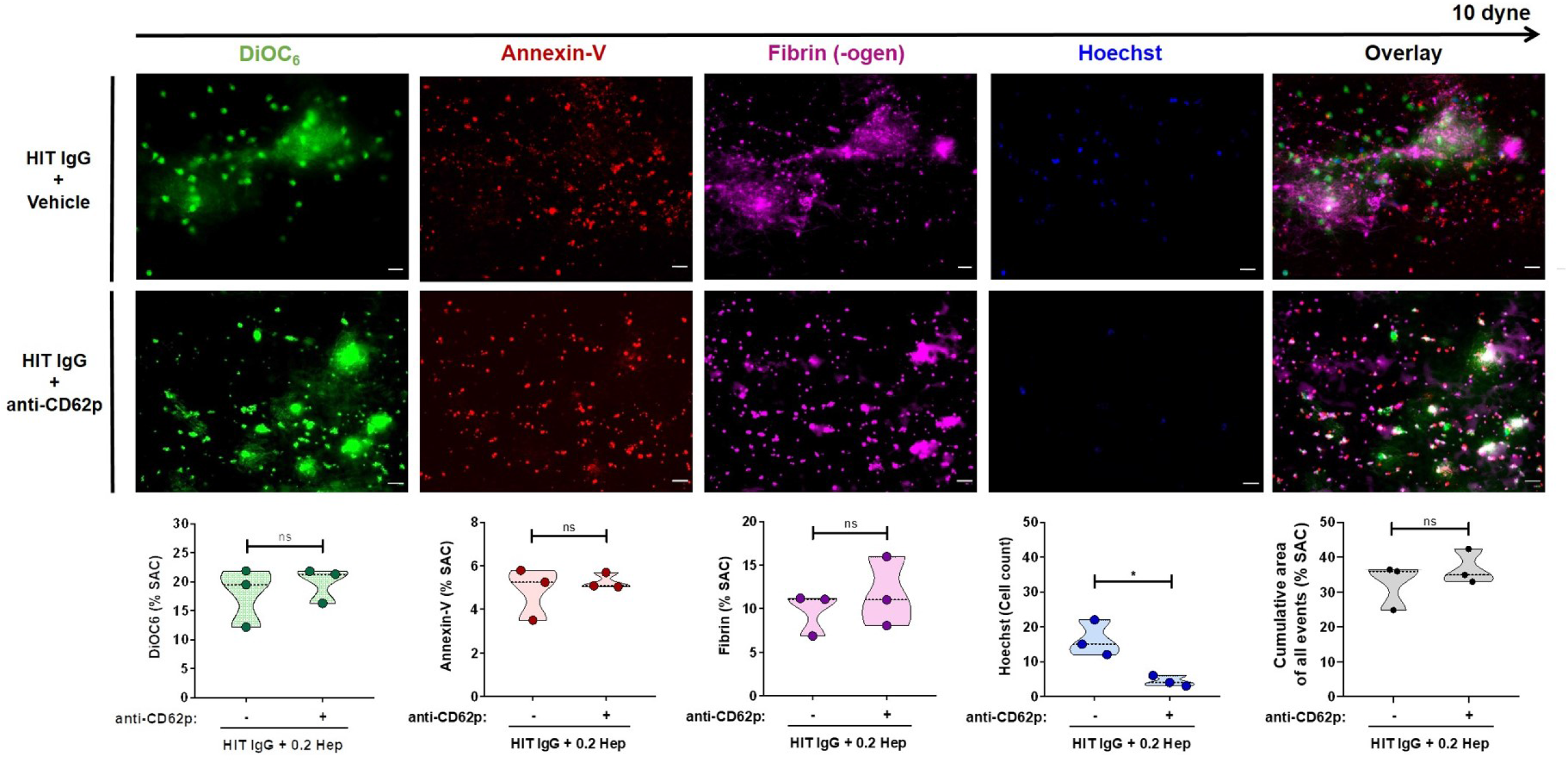
HIT antibodies can induce thrombus without platelet-leukocyte interaction. PLTs from healthy individuals were incubated with HIT patient IgGs and treated with vehicle (upper panel) or anti-CD62p blocking Ab (lower panel). After reconstitution into autologous whole blood and recalcification, samples were perfused through microfluidic channels at a venous shear rate of 250s^-1^ (10 dyne) for 10 min. After perfusion, images were acquired at x40 magnification. Scale bar 20μm. Violin plots showing the percentage of total surface area coverage (%SAC) by DiOC_6_, PS, Fibrin (-ogen), count of Hoechst positive labelled cells and cumulative area of DiOC_6_, PS and Fibrin (-ogen) labelled thrombus in the microfluidic channel. *p<0.05, **p<0.01 and ***p<0.001. ns, non-significant; PLT, platelet; IgG, immunoglobulin G; PS, phosphatidylserine.

Finally, we investigated the prothrombotic role of PS on the surface of HIT Ab-induced procoagulant PLTs. Notably, inhibition of procoagulant PS with Lactadherin resulted in a significant inhibition of HIT Ab-induced procoagulant PLT-mediated thrombin generation compared to vehicle (mean peak thrombin [nM]±SEM: 305.60±22.24 vs. 151.80±30.28, p value 0.0040; Figure 5C+D). Our hypothesis that increased PS could be the critical mediator of increased thrombus formation was further reinforced as PS blockade with Lactadherin resulted in significant reduction of HIT Ab-induced thrombus formation. Treatment of PLTs with Lactadherin upon HIT IgG incubation resulted in marked reduction of procoagulant PLT deposition (mean %SAC Annexin-V±SEM: 4.29±0.82% vs. 0.912±0.20%, p value 0.0357; Figure 7). Additionally, the presence of Lactadherin also prevented non-procoagulant PLT deposition on the collagen surface (mean %SAC DiOC_6_±SEM: 18.25±2.58% vs. 2.20±0.58%, p value 0.0080). Most importantly, PS inhibition interfered with plasmatic coagulation as a nearly complete inhibition of fibrin network generation was observed (mean %SAC Fibrin(-ogen)±SEM: 10.83±0.53% vs. 2.88±1.30%, p value 0.0067). Interestingly, also multicellular thrombus composition was affected, as a reduction of leukocytes, although not significant, was observed in the presence of Lactadherin (mean no. of Hoechst positive cells±SEM: 14±6 vs. 1±1, p value 0.0602; Figure 7).

**Figure 7.**
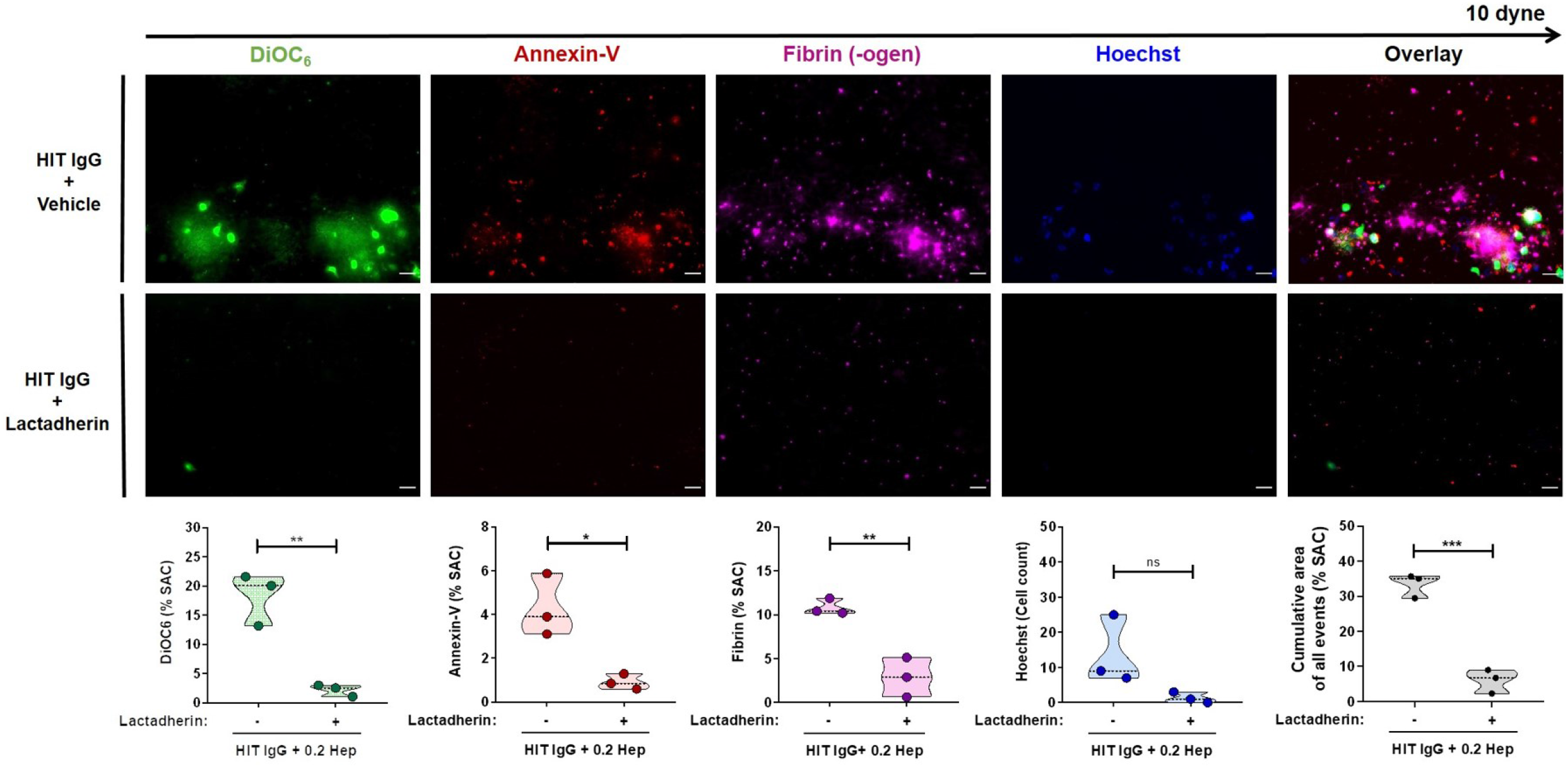
Platelet phosphatidylserine externalization is essential for HIT Ab-induced thrombus formation. PLTs from healthy individuals were incubated with HIT patient IgGs and treated with vehicle (upper panel) or Lactadherin (lower panel). After reconstitution into autologous whole blood and recalcification, samples were perfused through microfluidic channels at a venous shear rate of 250s^-1^ (10 dyne) for 10 min. Images were acquired at x40 magnification. Scale bar 20μm. Violin plots showing the percentage of total surface area coverage (%SAC) by DiOC_6_, PS, Fibrin (-ogen), count of Hoechst positive labelled cells and cumulative area with DiOC_6_, PS and Fibrin (-ogen) positive labelled thrombus in the microfluidic channel. *p<0.05, **p<0.01 and ***p<0.001. ns, non-significant; PLT, platelet; IgG, immunoglobulin G; PS, phosphatidylserine.

## Discussion

Thromboembolic events leading to high morbidity and mortality are frequent and still unpredictable complications in HIT.^3^ Of all laboratory findings, thrombocytopenia seems to be the most pronounced change associated with thrombosis in HIT.^9,10^ Although recent studies indicate the involvement of multicellular effector mechanisms, the question whether PLTs are capable of initating a prothrombotic response in HIT, remains elusive. In this study, we found that HIT IgG Abs isolated from serologically and clinically confirmed HIT cases induce a procoagulant PLT subpopulation. This was confirmed by the ability of procoagulant PLTs to cause increased thrombin generation and thrombus formation in a heparin-dependent fashion. Our data indicate that Fc-gamma-RIIA and subsequent intracellular calcium-dependent signaling pathways mediate the generation of procoagulant PLTs in HIT. Most importantly, we found that procoagulant PLT-induced prothrombotic conditions are mediated by externalized PS rather than P-selectin. These findings direct towards an essential role of procoagulant PLTs in the pathophysiology of thromboembolic complications in HIT. Our findings might have potential diagnostic as well as therapeutic relevance for the management of patients with suspected HIT.

We observed that the ability to generate procoagulant PLTs is restricted to sera from patients with laboratory and clinically confirmed HIT. Procoagulant PLT-formation was heparin-dependent indicating that the formation of PF4/heparin-HIT Ab immune complexes might be the underlying cause for procoagulant PLT-formation. In fact, IgG fractions from these sera modulate the expression of P-Selectin (CD62p) and PS on the surface of PLTs. Hence, the ability of HIT immune complexes to induce a procoagulant PLT phenotype seems to be independent of costimulatory signals i.e. additional PLT activation by extracellular agonists like thrombin. This observation contradicts recent findings, that report dual receptor stimulation as a requirement for procoagulant PLT formation.^6,36^ The discrepancy could be due to different experimental settings. We incubated isolated PLTs with HIT Abs in the absence of other cell types throughout our experiments.^37^ A higher concentration of PF4/heparin-HIT Ab immune complexes in the fluid phase could interact directly with PLT Fc-gamma-RIIA in our experimental conditions. This interaction could amplify Fc-gamma-RIIA-mediated signaling events and result in increased procoagulant PLT formation, despite the lack of other costimulatory signals.

To evaluate the biological relevance of HIT Ab-induced procoagulant PLTs, we analyzed the interaction with the plasmatic coagulation system and other blood cells using a PLT-based thrombin generation assay and an ex vivo thrombosis model, respectively. HIT Ab-induced procoagulant PLTs mediated significant increase in thrombin generation in the presence of heparin indicating a potential capability of causing ex vivo thrombosis. We optimized our thrombosis model to allow specific analysis of PLT-mediated thrombosis. In this model, PLTs that were incubated with IgGs from HIT patients but not HCs, induced increased thrombus formation in the presence of heparin at venous shear rates. Interestingly, PLT aggregates showed two different PLT phenotypes, namely procoagulant PLTs (Annexin-V positive/DiOC_6_ negative) and non-procoagulant PLTs (Annexin-V negative/DiOC_6_ positive).^38,39^ Recent evidence highlights that procoagulant PLTs primarily evolve on the surface of the growing thrombus where they promote thrombin generation.^40^ In our model, circulating HIT Ab-induced procoagulant PLTs attached to the collagen surface and later incorporated directly into the growing thrombus. Apart from this PS positive PLT aggregates, non-procoagulant PLTs also adhered to collagen in our model. The fact that the recruitment of non-procoagulant PLTs was inihibited by PS blockade suggests that procoagulant PLTs are not only an important component in the thrombus structure but also propagate the deposition of other PLT subpopulations as well as immune cells into the growing thrombus body.^41^

Of potential therapeutic interest is the observation that the prostacyclin analogue Iloprost significantly reduces HIT Ab-induced procoagulant PLT and thrombus formation. In fact, in a recent clinical study patients with acute HIT undergoing cardiac surgery were treated with Iloprost in addition to heparin.^42^ HIT Ab levels remained stable and no HIT Ab-induced reactivity on PLTs was observed during the procedure, despite the presence of heparin. Compared with a non-HIT control group, the incidence of thrombotic and theromboembolic events was similar (5.4% vs 5.1%, respectively). Nevertheless, according to current HIT therapy guidelines, the use of prostacyclins is limited to cardiac surgery patients with acute HIT where treatment delay is not feasible.^23^ Our results suggest that Iloprost may have a potential role in the treatment of a broader HIT patient collective. However, further clinical studies are needed to test the effectiveness and safety of Iloprost in patients with HIT.

HIT Ab-induced PLT-neutrophil interplay via P-Selectin/P-Selectin glycoprotein ligand-1 was reported to result in NET formation and subsequent increased thrombosis in mice.^16,43^ In our experiments, blocking P-Selectin reduced the recruitment of leukocytes to the thrombus, but did not affect procoagulant PLT-mediated thrombin generation or thrombus formation. This finding indicates that HIT Ab-induced procoagulant PLTs can cause thrombus formation in the absence of direct PLT-leukocyte interactions. In contrast, the blockade of PS with Lactadherin significantly impaired the ability of HIT Ab-induced procoagulant PLTs to promote thrombin generation and thrombus formation. These data indicate an indispensable role of PS on HIT Ab-induced procoagulant PLTs to link cellular with plasmatic components of the coagulation cascade, leading to thrombin burst and subsequent fibrin network formation. Lactadherin also prevented the deposition of DiOC_6_ positive labelled PLTs in the thrombus structure. This observation enforces our theory that procoagulant PLTs might result in the deposition of other PLT phenotypes such as activated PLTs.

Findings of our study might have several important clinical implications. Currently no predictive biomarker exists that distinguish between HIT patients with high risk for thrombosis and those with ‘‘only’’ thrombocytopenia. The detection of procoagulant PLTs in HIT patients via FC could help in risk stratification for prophylactic or therapeutic anticoagulation. Without any doubt, targeting PS to inhibit propagation of the prothrombotic condition in HIT is another promising clinical aspect of our results.

This study has limitations. First, our data are based on experiments using sera from a relatively small number of patients. Second, our study does not provide data on the dynamics of thrombus formation. Future research attempts should focus on the sequence of events and the relevance of intercellular interactions to enable better identification of procoagulant PLTs role in the initiation and propagation of thrombus formation. Finally, while our ex vivo thrombosis model might provide information on the interaction between blood cells, further investigations are needed to assess the influence of other cell types (e.g. endothelial cells).

In conclusion, our study suggests an indispensable role of procoagulant PLTs in the pathophysiology of HIT-associated thrombosis. The inhibition of HIT Ab-induced procoagulant PLT formation via Iloprost or the inhibition of prothrombotic effects with PS targeting specific therapeutics could be a promising approach to prevent the onset of thromboembolic events in HIT patients without increasing the risk for anticoagulant-related bleeding.

## Acknowledgements

We thank Flavianna Rigoni and Andreas Witzemann for their excellent technical support. The authors thank Susanne Staub for editing the article as native English speaker.

This work was supported by grants to T.B. from German Red Cross and the Herzstiftung to T.B. (BA5158/4 and TSG-Study). This project was supported by the German Research Foundation (DFG) – Project number 374031971 – TRR 240.

## Author contributions

J.Z. and T.B. designed the study. K.A and T.B. were responsible for the treatment of the patients. J.Z. and K.A. collected and analyzed the clinical data. J.Z., A.S., K.W, H.J., and K.A. performed the experiments. J.Z., A.S., K.A., G.U and T.B. analyzed the data, interpreted the results and wrote the manuscript. All authors read and approved the manuscript.

