## Supplemental Material for "Platelet phosphatidylserine is the critical mediator of thrombosis in heparin-induced thrombocytopenia"

### Supplemental Methods

#### Testing for anti-PF4/heparin antibodies

A commercially available IgG-EIA was used in accordance to manufacturer's instructions (Hyphen Biomed, Neuville-sur-Oise, France). A sample was considered reactive if the optical density (OD) was higher than 0.500. The ability of sera to activate PLTs was tested using the HIPA as previously described.[19] In brief, serum was tested with washed platelets (wPLTs) from four different healthy donors in the absence (buffer alone) or in the presence of heparin (0.2 IU/mL and 100 IU/mL). Reactions were placed in microtiter wells containing spherical stir bars and stirred at approximately 500 revolutions per minute (rpm). Wells were examined optically at five-minute (min) intervals for loss of turbidity. A serum was considered reactive (positive) if a shift from turbidity to transparency occurred within 30 min in at least two PLT suspensions. Observation time was 45 min. Each test included a diluted serum from a patient with HIT as a weak positive control, collagen (5µg/mL) as strong positive control and a serum from a healthy donor as a negative control.

#### Preparation of washed platelets

wPLTs were prepared from venous blood samples as previously described.[19,20] Briefly, fresh whole blood (WB) from healthy donors was withdrawn by cubital venipuncture into acidic-citrate-dextrose (ACD) containing vacutainers (Becton-Dickinson, Plymouth, United-Kingdom) and allowed to rest for 45 min at 37°C. After a centrifugation step (120g, 20 min, room temperature [RT], no brake), PLT-rich-plasma (PRP) was gently separated and supplemented with apyrase (5 µL/mL; Sigma-Aldrich, St. Louis, USA) and pre-warmed ACD (333 µL/mL; Sigma-Aldrich, St. Louis, USA).

After an additional centrifugation step (650g, 7 min, RT, no brake), the PLT pellet was resuspended in 5 mL of wash-solution (modified Tyrode buffer: 5 mL bicarbonate buffer, 20 percent [%] bovine serum albumin, 10% glucose solution [Braun, Melsungen, Germany], 2.5 U/mL apyrase, 1 U/µL hirudin [Pentapharm, Basel, Swiss], pH 6.3) and allowed to rest for 15 min at 37°C. Following final centrifugation (650g, 7 min, RT, no brake) wPLTs were resuspended in 2 mL of resuspension-buffer (50 mL of modified Tyrode buffer, 0.5 mL of 1 mM MgCl2, 1 mL of 2 mM CaCl2, pH 7.2) and adjusted to $300x10^3$ PLTs/µL after the measurement at a hematological analyzer (CELL-DYN Ruby, Abott, Wiesbaden, Germany) was performed.

**Immunoglobulin G preparation**

IgG fractions were isolated from HIT as well as from control sera using a commercially available IgG-purification-kit (Melon™-Gel IgG Spin Purification Kit, Thermo Fisher Scientific, Waltham, USA) as recommended by the manufacturer. In brief, heat inactivated serum was diluted 1:10 in purification buffer and incubated with the kit specific gel IgG Purification Support for 10 min. Subsequently centrifugation through a 10µm pore size filter tube was performed for 1 min at 5800g. The flow-through was collected into 100 kDa pore sized centrifugal filters (Amicon Ultra-4, Merck Millipore, Cork, Ireland) with subsequent concentration to the initial volume of the used serum sample via centrifugation (10-15 min, 2000g, 4°C, with brake). IgG concentrations were measured using NanoDrop One Spectrophotometer at an excitation wavelength of 340 nm (Thermo Fisher Scientific, Waltham, USA).

**Treatment of PLTs with sera/IgGs**

37.5 µL of wPLTs or PRP were supplemented with 5 µL serum/IgG from HIT patients or controls and incubated for 1 hour under rotating conditions at RT. Afterwards, 5µL of the PLT suspension containing ~1x$10^6$ PLTs were transferred into a final volume of 100 µL of Hank´s balanced salt solution (HBSS) containing 137 mM NaCl, 1.25 mM $CaCl_2$, 5.5 mM glucose (Carl-Roth, Karlsruhe, Germany). Cell suspension was then incubated with 1 µL anti-CD62p-APC (BD, San Jose, USA), 1 µL Annexin-V-FITC (Immunotools, Friesoythe, Germany) and 2 µL anti-CD42a-PerCP (BD, San Jose, USA) for 30 min at RT in the dark. PLTs that were treated with thrombin receptor activating peptide-6 (TRAP-6; 5 µM, 30 min at RT) and ionomycin ([5 µM, 15 min at RT], both from Sigma-Aldrich, St. Louis, USA) served as positive controls. Afterwards, PLTs were resuspended with HBSS to a final volume of 500 µL and immediately assessed via flow cytometry (FC; Navios, Beckman-Coulter, Brea, USA).

**Thrombin generation assay**

HIT Ab-induced thrombin generation (TG) on PRP was detected using Calibrated Automated Thrombogram (CAT; Stago, Maastricht, Netherlands) according to the manufacturer’s instructions. In brief, venous blood from healthy individuals was withdrawn into vacutainers containing sodium citrate 0.105 M (3.2%; BD, Plymouth, UK) and allowed to rest for 20 min at RT. PRP was prepared by centrifugation (20 min, 120g, no brake) and adjusted with autologous platelet poor plasma (PPP; 10 min, 2000g, RT) to a PLT count of 150x$10^6$/mL. Afterwards, PRP was treated with IgGs from HC or HIT patients in the presence of buffer or low- (0.2 IU/mL) dose heparin and incubated for 60 min at RT under rotating conditions. Following incubation, samples were washed (10 min, 700g, no brake) and the remaining PLT pellet gently resuspended with fresh PPP. 80 µl of the cell suspension were dispensed into the well of round-bottom 96 well-microtitre plates (Fluoroskan Ascent, ThermoLabsystems, Helsinki, Finland) and supplemented with 20 µL of PRP-reagent containing recombinant tissue factor and low amounts of phospholipids (Thrombinoscope BV, Maastricht, The Netherlands). 20 µL of fluorogenic substrate and calcium (FluCa-Kit reagent, Thrombinoscope BV, Maastricht, The Netherlands) were dispensed automatically by the device in each well. Fluorescence was acquired for 60 min with a 390-nm excitation/460-nm emission filter set, and parameters automatically calculated by dedicated software (Thrombinoscope BV, Maastricht, The Netherlands). When indicated, PLTs were preincubated with moAb IV.3 or isotype control (20 µg/mL) for 30 min at RT. To investigate the effect of cAMP elevation on PLTs thrombin generation potential, PRP was preincubated with Iloprost (20nM) or vehicle for 5 min at RT prior to the incubation of HIT IgGs. For specific inhibition of CD62p and PS, PLTs were incubated with indicated concentrations of anti-CD62p and Lactadherin (both 15 min at RT) after HIT IgG incubation and further handled as described above.

### Supplemental Figures

#### Supplemental Figure 1

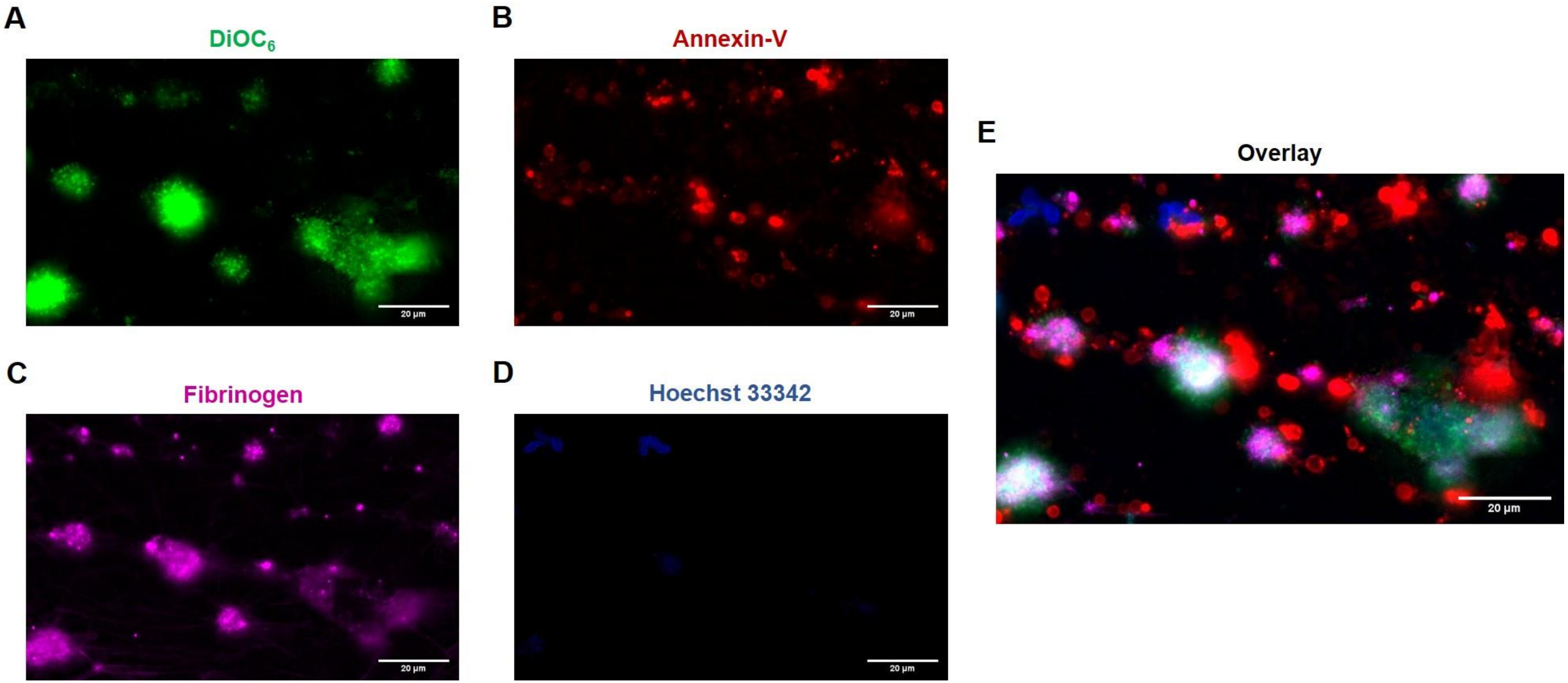


**Supplemental Figure 1-Phenotype of HIT antibody-induced multicellular thrombus**. Representative images of thrombus induced by PLTs after incubation with HIT IgG in the presence of low-dose (0.2 IU/mL) heparin. During reconstitution into autologous WB, samples were labelled with $DiOC_6$ (green) to identify non-procoagulant PLTs [**A**], AF647 Annexin-V (red) to detect procoagulant PLTs

[**B**] and AF546 human-Fibrinogen (magenta) and Hoechst 33342 (blue) for the detection of fibrin deposition [**C**] and leukocyte [**D**] recruitment, respectively. [**E**] shows overlay of all image sections. After 10 min perfusion at venous shear rates ($250s^{-1}$ [10 dyne]), images were acquired at x40 magnification. Scale bar 20µm. PLT, platelet; IgG, immunoglobulin G; WB, whole blood.

**Supplemental Figure 2**

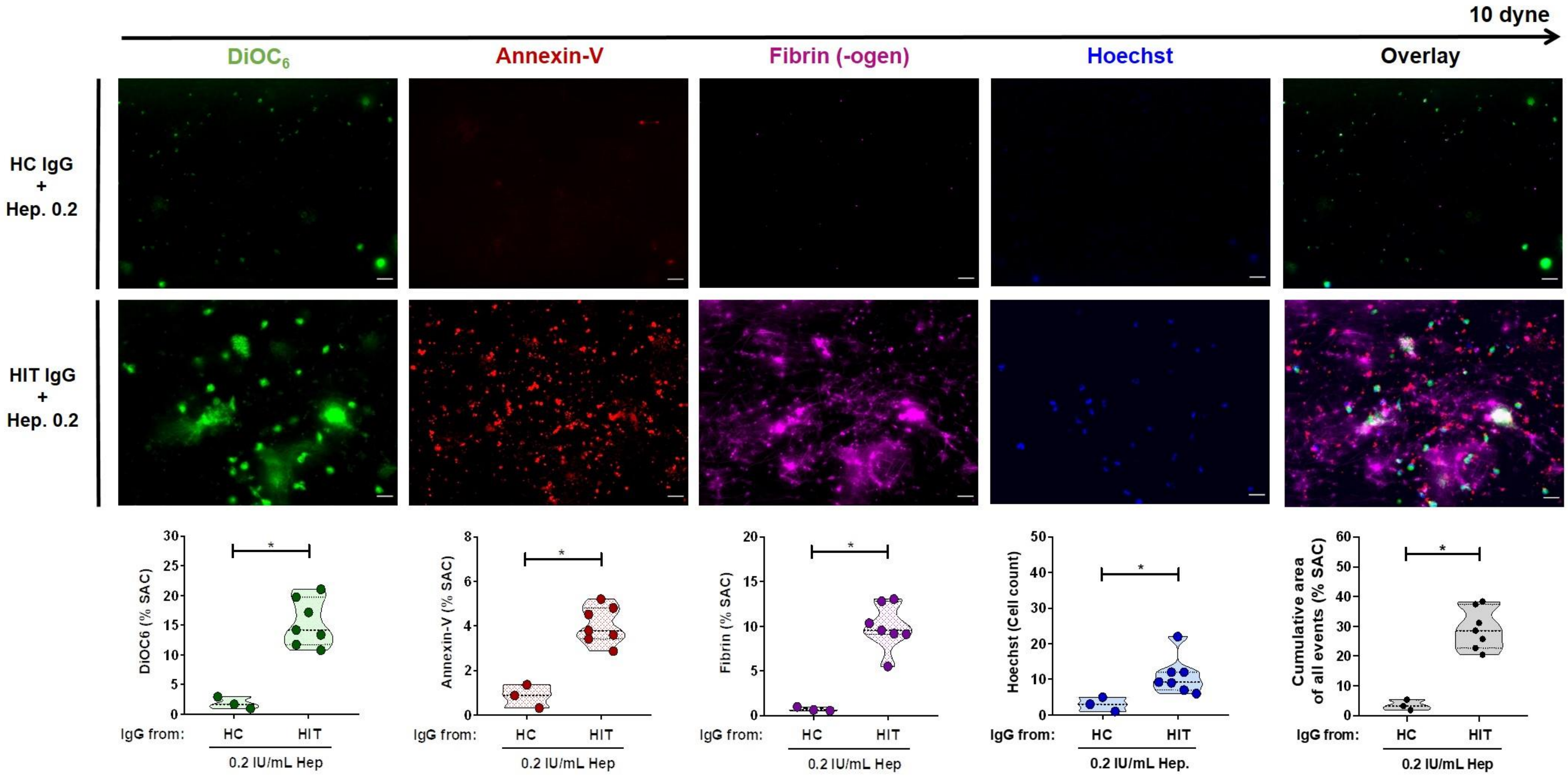


**Supplemental Figure 2-Procoagulant platelets contribute to anti-PF4/heparin IgG mediated thrombus formation.** PLTs from healthy individuals were incubated with IgGs from controls (HC) or HIT patients in the presence of low-dose (0.2 IU/mL) heparin and labelled with $DiOC_6$, AF647 Annexin-V, AF546 Fibrinogen and Hoechst 33342 prior to reconstitution into autologous whole blood and perfusion through microfluidic channels. After perfusion, images were acquired at x40 magnification. Scale bar 20µm. Violin plots showing the percentage of total surface area coverage (%SAC) by $DiOC_6$, PS, Fibrin-(ogen), count of Hoechst positive labelled cells and cumulative area of $DiOC_6$, PS and Fibrin-(ogen) labelled thrombus in the microfluidic channel. *$p<0.05$, **$p<0.01$ and ***$p<0.001$. ns, non-significant; HC, healthy control; PLT, platelet; PF4, platelet factor 4; IgG, immunoglobulin G; PS, phosphatidylserine.

**Supplemental Figure 3**

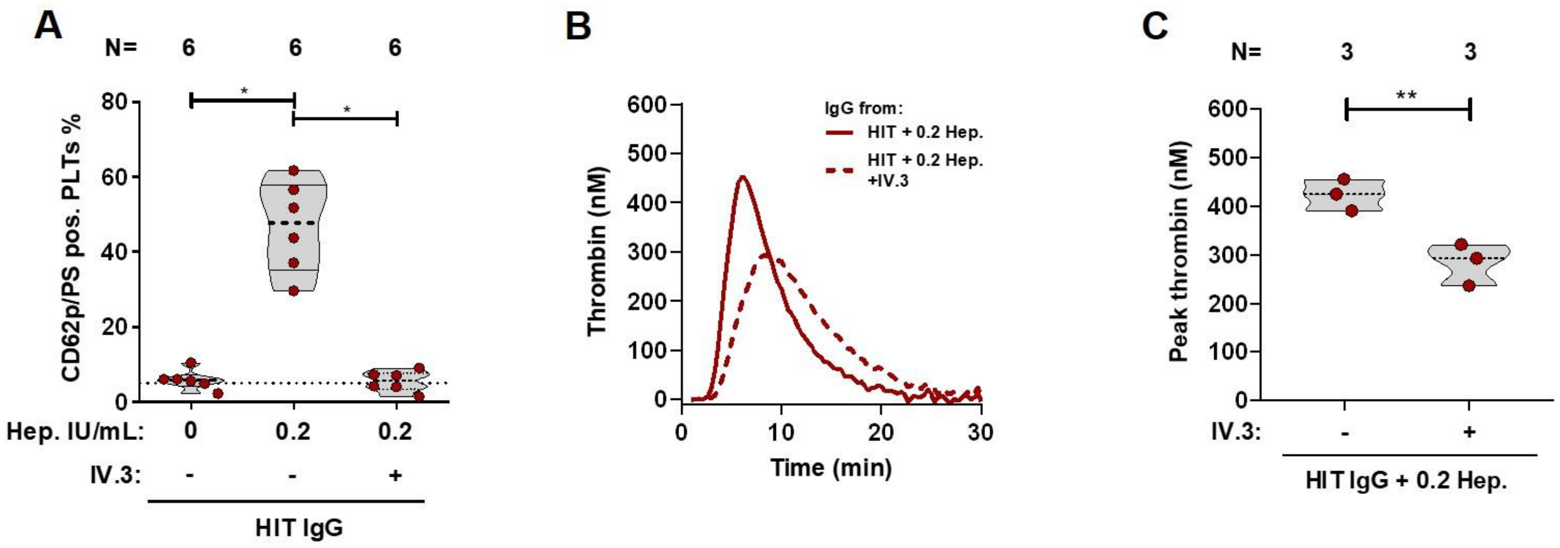


**Supplemental Figure 3-Inhibition of Fc-gamma-RIIA signal transduction prevents generation of procoagulant platelets and increased thrombin generation by HIT antibodies.** [**A**] HIT IgG-induced changes in PLTs PS externalization and CD62p expression were analyzed via Annexin-V-FITC and CD62p-APC double staining in the presence of isotype (-) or IV.3 moAb. Data are shown as percentage of Annexin-V-FITC and CD62p-APC double positive-labeled PLTs. [**B**] Representative thrombin generation curve induced on PLTs after incubation with IgG from different HIT patients in the presence of isotype (-) or IV.3 moAb. Each curve represents the amounts of generated thrombin over time induced by HIT IgG in the presence of heparin (0.2 IU/mL) and isotype (solid red line) or IV.3 (dashed red line) moAb. [**C**] Data were quantified as Peak thrombin generated (nM) using thrombinoscope software and graphpad prism.

*P < 0.05, **P < 0.01, ***P < 0.001, and ****P < 0.0001. CAT, Calibrated Automated Thrombogram; IgG, immunoglobulin G; moAb, monoclonal Ab, PLT, platelet; PRP, platelet-rich plasma; CD62p, P-selectin; PS, phosphatidylserine. N, number of samples.

**Supplemental Figure 4**

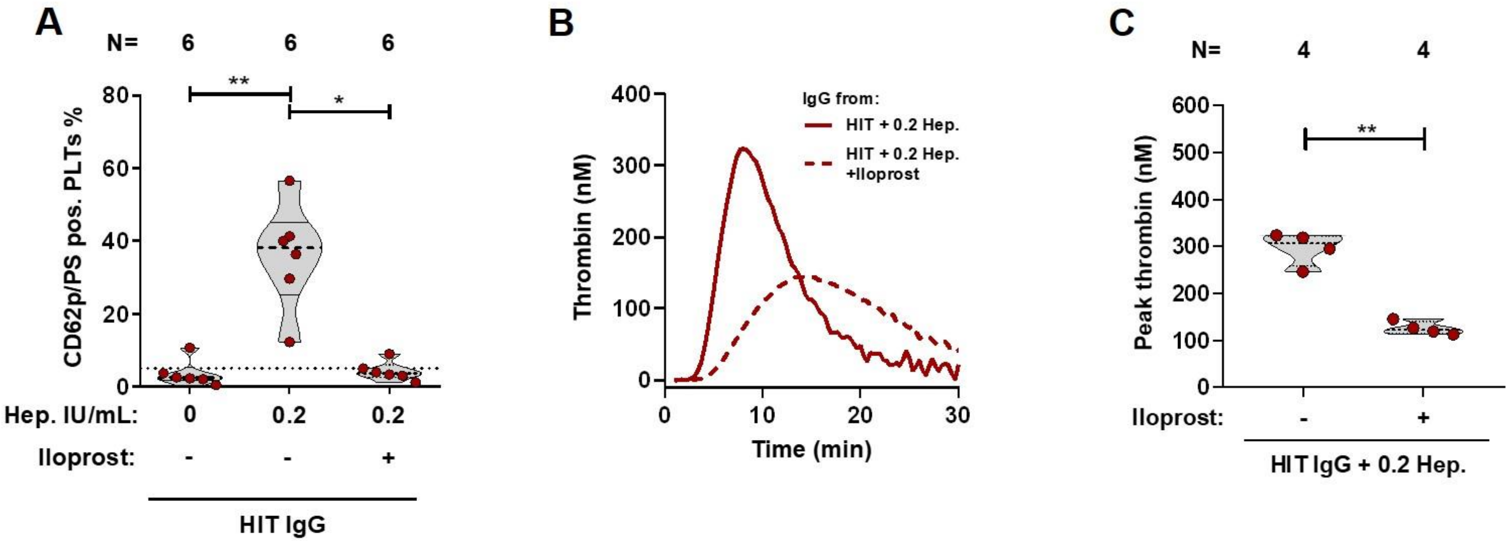


**Suppl. Figure 4-HIT antibody-induced procoagulant platelet and increased thrombin formation is calcium-dependent.** [**A**] HIT IgG-induced changes in PLTs PS externalization and CD62p expression were analyzed via Annexin-V-FITC and CD62p-APC double staining in the presence of vehicle (-) or Iloprost (20nM). Data are shown percentage±SEM of Annexin-V-FITC and CD62p-APC double positive-labeled PLTs. [**B**] Representative thrombin generation curve induced on PLTs after incubation with IgG from different HIT patients in the presence of heparin (0.2 IU/mL) and vehicle or Iloprost. Each curve represents the amounts of generated thrombin over time induced by HIT IgG in the presence of vehicle (solid red line) or Iloprost (dashed red line). [**C**] Data were quantified as Peak thrombin generated (nM) using thrombinoscope software and graphpad prism. *P < 0.05, **P < 0.01, ***P < 0.001, and ****P < 0.0001. CAT, Calibrated Automated Thrombogram; IgG, immunoglobulin G; PLT, platelet; PRP, platelet-rich plasma; CD62p, P-selectin; PS, phosphatidylserine; N, number of samples.
